# Natural Killer cells dampen the pathogenic features of recall responses to influenza infection

**DOI:** 10.1101/846626

**Authors:** Jason P. Mooney, Tedi Qendro, Marianne Keith, Adrian W. Philbey, Helen T. Groves, John S. Tregoning, Martin R. Goodier, Eleanor M. Riley

**Author notes:** Correspondence address: Jason P Mooney PhD, The Roslin Institute, University of Edinburgh, Easter Bush, Midlothian, EH25 9RG, United Kingdom.

## Abstract

Despite evidence of augmented Natural Killer (NK) cell responses after influenza vaccination, the role of these cells in vaccine-induced immunity remains unclear. Here, we hypothesized that NK cells might increase viral clearance but possibly at the expense of increased severity of pathology. On the contrary, we found that NK cells serve a homeostatic role during influenza virus infection of vaccinated mice, allowing viral clearance with minimal pathology. Using a diphtheria toxin receptor transgenic mouse model, we were able to specifically deplete NKp46+ NK cells through the administration of diphtheria toxin. Using this model, we assessed the effect of NK cell depletion prior to influenza challenge in vaccinated and unvaccinated mice. NK-depleted, vaccinated animals lost significantly more weight after viral challenge than vaccinated NK intact animals, indicating that NK cells ameliorate disease in vaccinated animals. However, there was also a significant reduction in viral load in NK-depleted, unvaccinated animals indicating that NK cells also constrain viral clearance. Depletion of NK cells after vaccination, but 21 days before infection, did not affect viral clearance or weight loss - indicating that it is the presence of NK cells during the infection itself that promotes homeostasis. Further work is needed to identify the mechanism(s) by which NK cells regulate adaptive immunity in influenza-vaccinated animals to allow efficient and effective virus control whilst simultaneously minimizing inflammation and pathology.

## Introduction

Influenza viruses are a significant cause of respiratory tract infections, leading to seasonal epidemics and unpredictable pandemics. Globally, influenza affects approximately 20-30% of children and 5-10% of adults, resulting in 3-5 million cases of severe illness with up to 500,000 deaths, annually (WHO, 2014). The very young, old, and immunocompromised are at greatest risk of succumbing to severe illness and death (Lewis, 2006). Currently, vaccination with live-attenuated or inactivated influenza virus is the most effective method for reducing infections at both the individual and population level (Cox and Subbarao, 1999;Ting et al., 2017).

Although adaptive immune responses play a key role in the resolution of influenza infection, innate immune responses play an essential role in restricting virus replication and thus limiting the scale of the initial infection (Iwasaki and Pillai, 2014). Amongst other innate immune players, Natural Killer (NK) cells can kill virus-infected cells and help to orchestrate nascent adaptive immune responses (White et al., 2008). Defects in NK cells are associated with increased susceptibility to viral infections (Biron et al., 1989;Orange, 2002). NK cell killing is mediated by cell surface cytotoxicity receptors (such as the natural cytotoxicity receptors NKp30, NKp44 and NKp46) binding to viral components or stress-induced ligands on the surface of virus-infected cells, leading to directed release of granzyme- and perforin-containing cytotoxic granules and/or Fas/Fas-ligand interactions (Smyth et al., 2005). Additionally, cytokine (particularly interferon (IFN)-γ) production by NK cells plays an integral role in shaping adaptive immunity (Vivier et al., 2008;Wagstaffe et al., 2018). For example, NK-derived IFN-γ can activate dendritic cell migration, thus promoting T cell priming (Ge et al., 2012). Evidence for the importance of NK cells in influenza virus infection comes primarily from studies in mice, although conclusions vary depending on the precise model employed (Table 1). For example, depletion of NK cells with anti-asialo GM1 antibody in mice and hamsters resulted in increased morbidity and mortality from influenza A virus infection (Stein-Streilein and Guffee, 1986). In addition, influenza infection is lethal in mice lacking the NK cell specific receptor NKp46 (NCR1) (Gazit et al., 2006), which has been reported to be a receptor for the influenza hemagglutinin (HA) protein (Arnon et al., 2001;Mandelboim et al., 2001). Conversely, others have reported that NK cell deficiency (whether through depletion with anti-asialo GM1 or anti-NK1.1, or due to a lack of interleukin (IL)-15) reduced weight loss and increased survival (Nakamura et al., 2010; Abdul-Careem et al., 2012;Zhou et al., 2013). Moreover, little is known regarding the role of NK cells in vaccine-induced immunity to influenza. We set out, therefore, to address the role of NK cells during acute influenza infection, before and after vaccination, using diphtheria-toxin (DT) mediated ablation of NK cells in genetically modified mice in which NKp46 expression drives expression of the DT receptor. Given that IL-2 produced from influenza-specific T cells is dependent on the NK cell driven IFN-γ response in early influenza infection (He et al., 2004;Wagstaffe et al., 2018), we hypothesized that NK cell ablation would impair viral clearance and increase disease severity.

**Table 1:** Methods of NK depletion during acute influenza infection.

| Author | Mouse Strain | Mouse Sex | Flu Strain | Dose of Flu Challenge | NK Depletion | Main Findings |
| --- | --- | --- | --- | --- | --- | --- |
| (Gazit et al., 2006) | 129/Sv, C57BL/6 | Not specified | A/PR/8/34 (H1N1) | $3 \times 10^3$ PFU (or 0.12 HAU), 60-70% survival in WT | NCR1-gfp knockout | Loss of NCR1 (NKp46) results in lethal infection (0% survival) |
| (Zhou et al., 2013) | C57BL/6 | Female | A/PR/8/34 (H1N1) | 5 HAU - 0% survival (in WT)<br>0.5 HAU - 30% survival (in WT)<br>0.0625 HAU - 80% survival (in WT) | Anti NK1.1* | High dose (5 HAU) infection + NK cell depletion improves survival.<br>Medium dose (0.5 HAU) infection + depletion decreases survival.<br>Low dose (0.0625 HAU) + depletion has no effect. |
| (Abdul-Careem et al., 2012) | C57BL/6 BALB/C | Male | A/PR/8/34 (H1N1) | $5 \times 10^3$ PFU (i.n.)<br>0% survival (in WT) | IL-15-/- KO mice, anti-NK1.1*, anti-asialo GM1<sup>†</sup> | NK cells depleted with anti-NK1.1 or anti-asialo GM1, and IL-15-/-KO mice had increased survival and lower weight loss compared to WT |
| (Nakamura et al., 2010) | C57BL/6 | Not specified | Mouse-adapted A/FM/1/47 (H1N1) | 1000 PFU (i.n.)<br>0% survival in WT | IL-15-/- KO mice<sup>‡</sup> | IL-15 deficiency improved survival of virus infected mice |
| (Stein-Streilein and Guffee, 1986) | B6D2F1 | Female | A/PR/8/34 (H1N1) | 0.125 HAU (intratracheal)<br>50% survival in WT mice. | Anti-asialo GM1<sup>†</sup> | NK cell depleted mice (and hamsters) had decreased survival to infection and increased viral titers |
| (Nogusa et al., 2008) | C57BL/6 | Male | A/PR/8/34 (H1N1) | $10^4$ TCID<sub>50</sub> HAU | Anti-NK1.1* | Mice depleted of NK cells lost significantly more weight than infected WT young mice and had greater viral loads. |
| (Kos and Engleman, 1996) | C57BL/6 | Female | A/WSN | 1000 HAU (i.p) | Anti-NK1.1* | NK cells were essential for the induction of influenza virus-specific CTL responses |

Note: Such methods of NK cell depletion have been shown not to be specific. \*NK1.1 is also used as a marker of NKT cells and
<sup>†</sup>anti-asialo GM1 can not only recognize T cells, but administration can also lead to basophil depletion (Nishikado et al., 2011).
<sup>‡</sup>IL-15 KO mice, while primarily exhibiting an NK cell deficiency, also suffer from T cell deficiencies and impaired lymphocyte trafficking (Lodolce et al., 1998; Itsumi et al., 2009). Therefore, we used NKp46 expression as the basis for NK cell depletion, as it has been shown to be expressed by mouse NK cells (Biaassoni et al., 1999), but also some type 1 innate lymphoid cells (Cortez and Colonna, 2016).

## Materials and Methods

### Ethics statement

All experiments were performed in accordance with United Kingdom (UK) Home Office Regulations under Project License 70/8291 and were approved by the animal welfare and ethical review board of the London School of Hygiene and Tropical Medicine (LSHTM).

### Mice

C57BL/6J and C57BL/6-Gt(ROSA)26Sor^tm1(HBEGF)Awai^/J mice were purchased from Jackson Laboratories via Charles River (Tranent, UK). C57BL/6-NKp46:iCre^+/+^ mice were kindly provided by Professor Eric Vivier (Centre d’Immunologie de Marseille-Luminy, Institut Universitaire de France, France). C57BL/6-Rosa26iDTR^+/+^ and C57BL/6-NKp46:iCre^+/+^ mice were crossed to generate F1 NKp46-iCre/Rosa26iDTR^+/−^ (referred to as NKp46-DTR). Groups of NKp46-DTR mice were age- and sex-matched for all experiments, with influenza challenge occurring between 8 and 10 weeks of age. Both sexes of the F1 generation were used in this study, with 68 female and 106 male NKp46-DTR mice being used in total. The sexes of mice used in each experiment are shown in the figure panels ({M} male, {F} female) and figure legends. All mice were maintained in individually ventilated cages in a specified pathogen free facility.

### Viral infection and vaccination

Egg grown influenza A/California/4/2009 virus was kindly provided by Dr. John McCauley (Crick Worldwide Influenza Centre, The Francis Crick Institute, London UK) at 128 hemagglutination units (HAU)/mL and stored at −80°C until use. For infections, mice were anesthetized by intraperitoneal (i.p.) injection of ketamine (100 mg/kg)/xylazine (10 mg/kg) and then inoculated intranasally (i.n.) with 30 μL of virus (0.5 HAU) diluted in Dulbecco’s phosphate-buffered saline (DPBS) (Gibco, Loughborough UK). Mock-treated control mice were inoculated similarly with DPBS. Mice were monitored and weighed daily to assess infection and euthanized at day 4 post-infection (p.i.), or if they reached the humane endpoint of 20% loss of body weight at an earlier time point. Twenty eight days prior to challenge, some mice were vaccinated by i.p. injection of the human 2015-2016 seasonal influenza vaccine (human Sanofi-Pasteur-MSD inactivated trivalent influenza vaccine (split-virion) containing influenza haemagglutinin (HA), including A/California/7/2009 (H1N1) pdm09-like strain NYMC X-179A), diluted 1:3 in DPBS with 500 μL per mouse with a final dose of 5 μg of each HA.

### Depletion of NK cells

For selective depletion of NK cells, NKp46-diphtheria toxin receptor (NKp46-DTR) mice were injected i.p. with 1.25 μg DT (diluted in 100 μL DPBS) on days 25 and 28 post-vaccination. Alternatively, mice were injected once with 2.5 μg DT and allowed to recover for 3 weeks prior to influenza challenge. Control (mock depleted) NKp46-DTR mice were injected i.p. with 100 μL DPBS.

### Isolation of lung cells and viral supernatant

Lungs were aseptically removed from euthanized mice and stored in 5 mL DPBS on ice until processing. Lung single-cell suspensions were obtained using methods previously described (Sauer et al., 2007). Briefly, isolated lungs were cut into 2-3 mm^2^ sections and resuspended in 5 mL digestion solution of Roswell Park Memorial Institute (RPMI) 1640 Medium (Gibco) supplemented with 5 mM GlutaMax (Gibco), 5% (v/v) fetal bovine serum (Gibco), and 1% (v/v) Penicillin-Streptomycin solution (10,000 units/mL, Thermo Fisher Scientific, Loughborough UK). Digestion solutions contained DNAseI (20 μg/mL) (Roche, Basel Switzerland), and LiberaseTM (0.2 digestion units/lung) (Roche). The lung suspensions were digested at 37°C for 60 minutes with gentle rotating and then processed on a gentleMACS dissociator (Miltenyi, Surrey UK). Lung homogenates were then sieved through a 40 μm nylon filter to remove debris (BD Biosciences, Berkshire UK) and sieve washed with 5 mL DPBS. After centrifugation, an aliquot of lung supernatant was frozen at −70°C for downstream <u>viral qPCR and IL-6 protein ELISA</u>. Cell pellets were then resuspended in 5 mL ACK Lysis Buffer (Lonza, Edinburgh UK), washed and resuspended with DPBS for flow cytometry.

### Flow cytometry

Cells were stained in 96-well U-bottom plates as described previously (Mooney et al., 2015). Briefly, cell suspensions were stained with antibodies to cell surface markers, fixed (Cytofix/Cytoperm; BD Biosciences) and stained for intracellular markers. Antibodies and dyes used for flow cytometry staining are shown in Table 2. Cells were acquired on an LSRII flow cytometer (BD Biosciences) using FACSDIVA software. Data were analyzed using FlowJo V10 (Tree Star). Gates were initially set on singlets, followed by leukocytes (removing high SSC-A and low FSC-A) and live cells. Subsequent gates were based on unstained and/or FMO controls.

**Table 2:** Antibodies and dyes used for flow cytometry

| Antigen | Clone |
| --- | --- |
| Zombie Aqua Dead/Live | Not Applicable |
| CD16.2 | 9e9 |
| NKp46 (CD335) | 29A1.4 |
| CD19 | 6D5 |
| CD3 | 145.2C11 |
| CD11b | M1/70 |
| NK1.1 | PK136 |
| CD69 | H1.2F3 |
| CD107a | 1D4B |
| Ly6C | HK1.4 |
| Ly6G | 1A8 |

### Viral burden determination

Influenza burden in lung-purified ribonucleic acid (RNA) was determined by one-step quantitative reverse transcription polymerase chain reaction (RT-qPCR) for the HA gene, as previously described (Lam et al., 2010). RNA was isolated from 500 μL of frozen lung cell supernatant (PureLink Viral RNA Mini Kit, Thermo). As a standard, RNA was isolated from stock virus (128 HAU/mL) and diluted 1:5 starting from 2.3 HAU equivalents per well. Purified RNA (5 μL of 25 μL elution) was mixed with 5 μL of TaqMan Fast Virus 1-step Master Mix (Thermo), 7 μL of ultra-pure water, and 1 μL each of forward primer SWH1-1080 (0.8 μM, GATGGTAGATGGATGGTACGGTTAT), reverse primer SWH1-1159 (0.8 μM, TTGTTAAGTAATYTCGTCAATGGCATT), and probe SWH1-1128 (0.25 μM, FAM-AGGATATGCAGCCGACCT-NFQMGB) with amplification conditions: 55°C, 30min; 95°C, 2min; 40 cycles of 95°C, 15 sec and 60°C, 30sec, on a 7900HT Fast RT-PCR system (Applied Biosystems, Loughborough UK). For viral burden from whole lung tissue, 500 ng of purified RNA (as reported below) was assayed in the same manner as above.

### RNA analysis from tissue

Whole lung tissue was stored in RNAlater (Ambion, Loughborough UK) at post-mortem examination. RNA was extracted from tissue using RNeasy extraction kit (Qiagen, Manchester UK) according to the manufacturer’s instructions, using a TissueRuptor probe (Qiagen) for tissue homogenization. RNA was then treated with DNAseI (Ambion) to remove genomic deoxyribonucleic acid (DNA) contamination. For a quantitative analysis of messenger RNA (mRNA) levels, 1□μg of total RNA from each sample was reverse transcribed in a 20 μL volume (SuperScript IV VILO Master Mix; Thermo), and 2□μL of complementary DNA (cDNA) was used for each real-time reaction. RT-qPCR was performed using the primers listed in Table 3, SYBR green (Applied Biosystems) and 7500 Real-Time PCR System (Applied Biosystems). Data were analyzed using the comparative threshold cycle (cT) method (Applied Biosystems). Target gene transcription of each sample was normalized to the respective levels of beta-Actin mRNA and represented as fold change over gene expression in control animals.

**Table 3:**
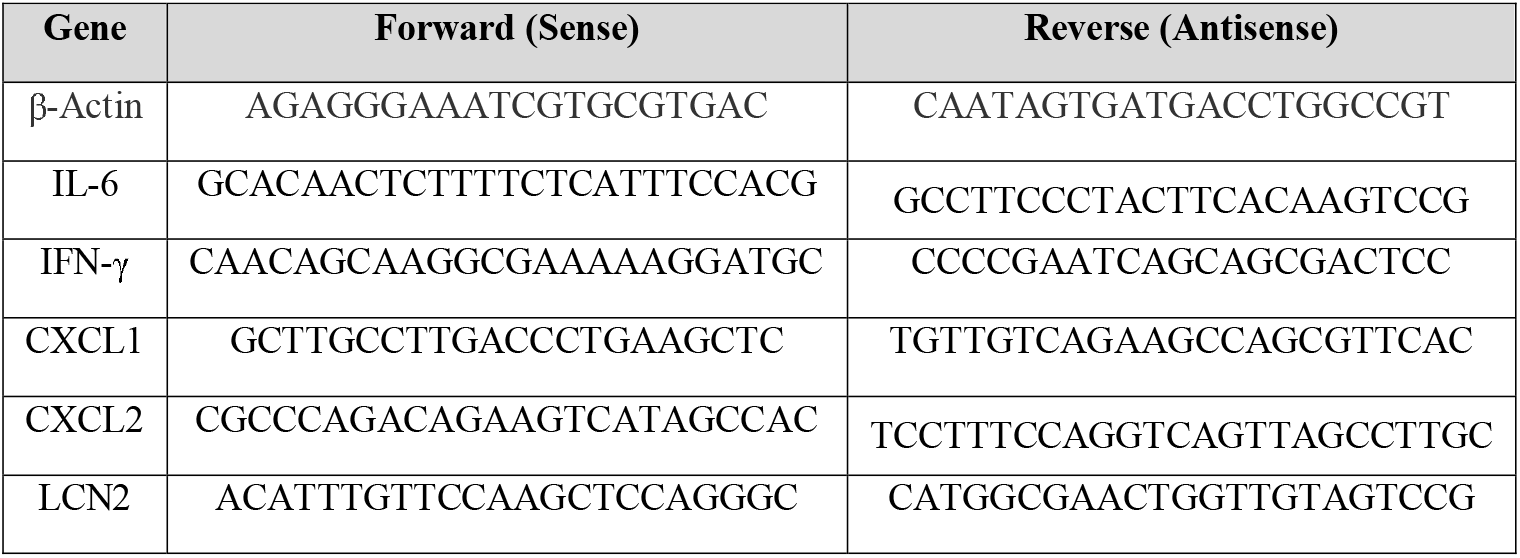
Primers used for lung qPCR

### Plasma analysis

Plasma was isolated by centrifugation of whole blood taken via cardiac puncture into a heparinized syringe and stored at −70°C. Plasma IL-6 levels were determined by sandwich enzyme-linked immunosorbent assay (ELISA) (Biolegend ELISA MAX Deluxe, London UK). Values below the blank were reported at the limit of detection for statistical purposes. To determine circulating influenza HA antibodies following influenza vaccination, a direct ELISA was done. Here, flat MaxiSorp 96-well plates (Nunc) were coated overnight with the vaccine (diluted 1:3000 in DPBS, or 0.01 μg/mL HA) or live virus (1.28 HAU/mL) at 100 μL. Using a commercial ELISA reagent kit (eBioscience catalogue number BMS412, Loughborough UK), plates were washed three times and blocked for 1 hour with 200 μL of assay buffer. After washing, 100 μL of serially-diluted plasma was incubated at room temperature (RT) for 2 hours. After washing, 100 μL of sheep, anti-mouse immunoglobulin (Ig)-G horseradish peroxidase (HRP)-linked secondary antibody (Amersham catalogue number NA931, GE Healthcare, Buckinghamshire UK) diluted 1:4000 was incubated at RT for 2 hours before final washes and 3,3’,5,5’-Tetramethylbenzidine (TMB) substrate development. Absorbance was determined at 450 nm by SpectraMax M5 microplate reader (Molecular Devices, Wokingham UK). Vaccine take was confirmed for all vaccinated mice with HA-IgG ELISA at 1:100 dilution of plasma.

### Histopathology

Sections were cut at 4 μm thickness from formalin-fixed, paraffin-embedded whole lung tissues and stained with hematoxylin and eosin (H&E) by the University of Edinburgh Easter Bush Pathology Laboratory and scored, in a blinded fashion, for inflammation (vasculitis, bronchiolitis, and alveolitis), oedema (perivascular oedema, peribronchiolar oedema, and alveolar oedema), leukocyte infiltration (perivascular, peribronchiolar, and in alveolar walls) and neutrophil infiltration (perivascular, peribronchiolar, and in alveolar walls). Each criterion was scored numerically as unremarkable (0), mild (1), mild to moderate (2), moderate (3), moderate to marked (4), and marked (5). Photomicrographs of histological sections were taken using a 20x objective with an Olympus BX41 microscope using an Olympus DP72 camera.

### Statistical analysis

Differences between groups were analysed by Mann-Whitney U test on combined data from two independent experiments, unless otherwise noted in legends. Spearman’s correlation was used to identify statistically significant associations between weight loss, influenza burden and lung neutrophils. All analyses were conducted using GraphPad Prism 7.

## Results

### Influenza vaccination reduces weight loss and viral burden in mice

To characterize the role of NK cells in influenza infection and immunization, we established a model of acute influenza infection in C57BL/6J mice (Fig. 1). C57BL/6J mice were infected i.n. with 0.5 HAU of influenza strain A/California/04/2009. Infected mice developed an acute infection, losing 20% of their body weight by 4 days post-infection (Fig. 1A). Mice were also vaccinated, intraperitoneally with the human Sanofi-Pasteur-MSD inactivated trivalent influenza vaccine (split-virion), four weeks prior to influenza challenge. Vaccinated mice lost significantly less weight (Fig. 1A) and had lower viral burden in their lungs (Fig. 1B) compared to unvaccinated mice; the reduction in influenza burden in the lung correlated with reduced weight loss (Fig. 1C).

**Figure 1.**
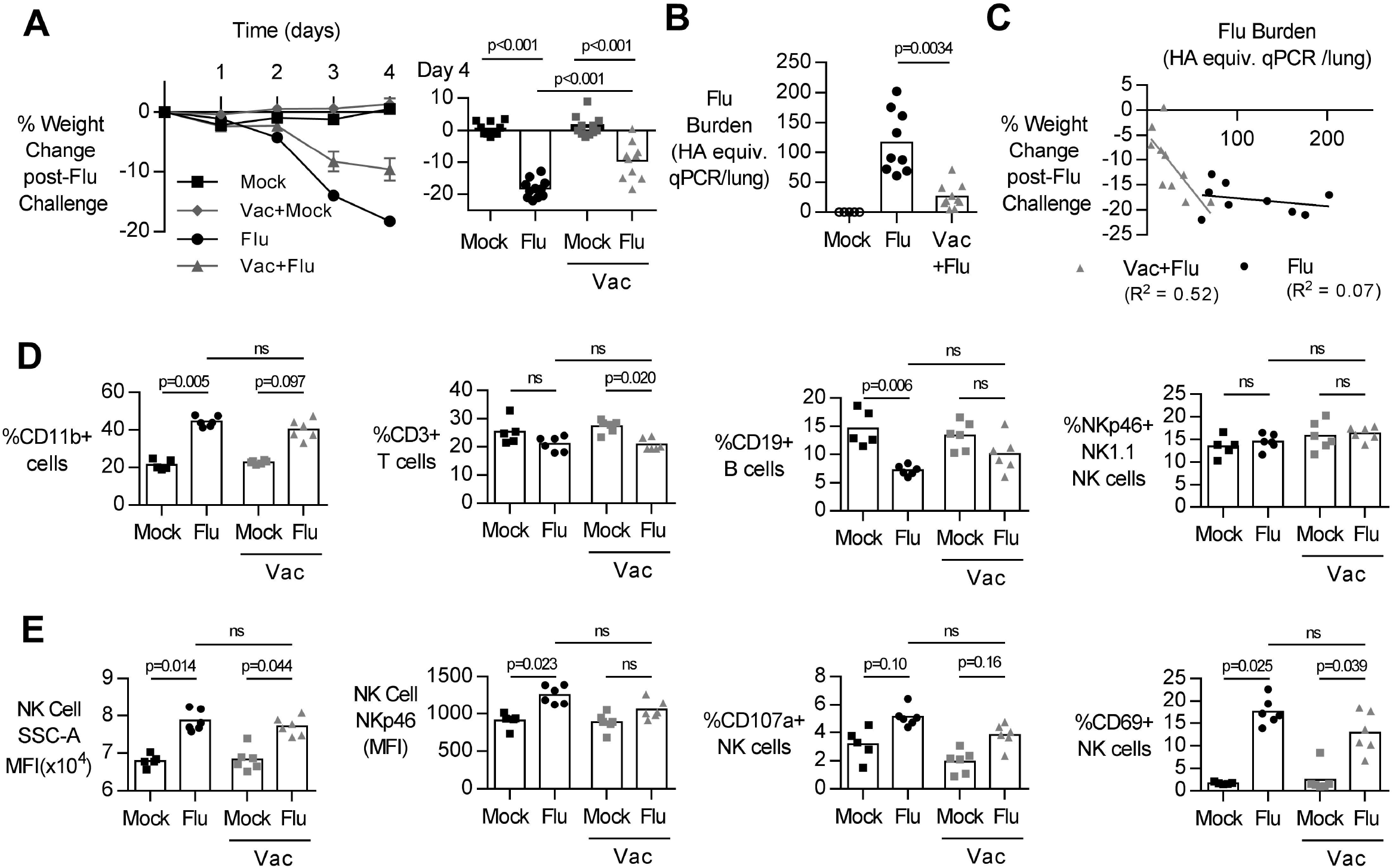
Primary influenza infection induces rapid weight loss and NK cell activation in lung but vaccination reduces weight loss and lung viral burden. C57BL/6 female mice were challenged intranasally with 5 hemagglutination units (HAU) of influenza A/California/4/2009 (Flu) or mock treated with DPBS (Mock). Four weeks prior to challenge, mice were vaccinated intraperitoneally with the trivalent Sanofi influenza vaccine (Vac). **(A)** Weight loss over 4 days. **(B-E)** At day 4 post infection, lungs were excised and cell-free supernatant was analyzed by qPCR for influenza viral burden (plotted against a dose curve of Flu with known HAU, giving HAU equivalents) **(B)** and plotted against weight loss **(C)**. Data fitted to a non-linear regression line with R square value shown **(C)**. **(D-E)** Lung cell pellets were analyzed by flow cytometry for **(D)** cellular abundance and **(E)** Natural Killer (NK) activation markers. Weight loss and viral burden data are a pool of two independent experiments (n=8-9/group), while flow data is one experiment (n=5/group). Dots represent individual mice with bars showing mean. Line data shown as mean±SEM. Significance determined by Mann-Whitney U test.

Acute influenza infection was marked by a significant increase in the frequency of CD11b+ cells in the lung, with a decrease in proportions of T (%CD3+) and B cells (%CD19+) (Fig. 1D). The frequency of NK cells, identified by NK1.1 and NKp46 co-expression (Walzer et al., 2007;Zhou et al., 2013), did not change after influenza infection (Fig 1D). However, there was a significant increase in the side-scatter (SSC) of the NK cell population (Fig. 1E), indicating increased activation (Brady et al., 2004;Skak et al., 2008;Marçais et al., 2014) and expression of NKp46, an activating NK cell receptor reported to bind influenza-derived HA (Gazit et al., 2006;Achdout et al., 2010;Glasner et al., 2012), was also increased in infected animals. Lastly, there was a significant increase in the frequency of NK cell activation marker CD69 (Fig. 1E). Together, these results suggest that the immune response to acute influenza infection, whether in a naïve or vaccinated animal, is characterized by an influx of CD11b+ cells and activated NK cells.

### Depletion of NK cells prior to influenza challenge reduces lung virus burden but increases weight loss in vaccinated mice

To evaluate the contribution of NK cells to vaccine-induced protection from influenza infection, transgenic mice (NKp46-DTR) were selectively depleted of NK cells by injection of DT (Fig. 2A) three days before infection. DT treatment resulted in a robust depletion of NK cells from the lung (Fig. 2B) but did not change circulating antibodies established by vaccination (Fig. 2C).

**Figure 2.**
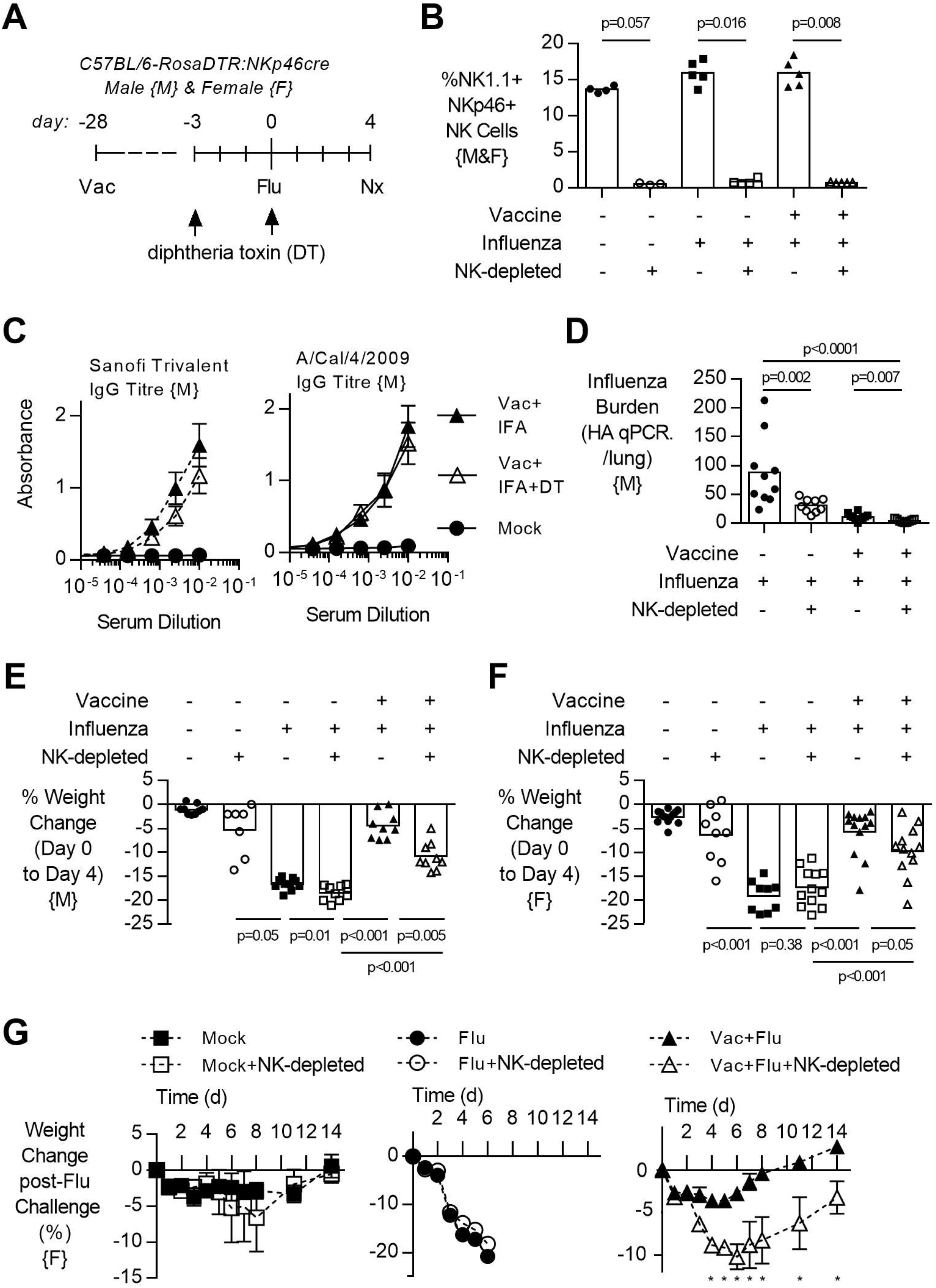
NK cell depletion reduces lung viral burden and increases weight loss in vaccinated, influenza-challenged mice. Transgenic C57BL/6 mice with NKp46 driven expression of diphtheria toxin (DT) receptor were vaccinated 28 days prior to intranasal influenza (flu) challenge, as in Fig. 1. **(A)** Immediately prior to infection, a subset of mice received two intraperitoneal injections of DT (1.25 μg). **(B)** Levels of NK1.1+, NKp46+ NK cells in the lung, as a proportion of singlet, live leukocytes at 4 days post infection (necropsy, nx). **(C)** Circulating IgG antibodies to both the vaccine and challenge virus with or without DT treatment at 4 days post infection. **(D)** Lung cell-free supernatants were analyzed by qPCR for influenza viral burden (plotted against a dose curve of IFA with known HAU, giving HAU equivalents). **(E-F)** Weight loss at day 4 in Male (E) and female (F) mice. **(G)** Weight loss followed for 14 days post challenge. (D-E) A pool of two independent experiments (n=8-9/group), while (B-C) is one experiment (n=5/group); all mice were male {M}. (F) A pool of three experiments (n=9-13/group) and (G) one experiment (n=4-5/group); all mice were female {F}. Dots represent individual mice with bars showing mean. Line data shown as mean±SEM. Significance determined by Mann-Whitney U test, ns = not significant. * represents p<0.05.

Depletion of NK cells prior to influenza challenge infection led to a significant decrease in influenza burden in the lung, regardless of vaccination status (Fig. 2D). Viral burden was 2.9 fold higher in naïve NK cell-intact mice than in naïve NK cell-depleted mice (p=0.002) and 2.6 fold higher in vaccinated NK cell-intact mice than in vaccinated NK cell-depleted mice (p=0.007) (Fig. 2D). However, infected NK-depleted animals lost more weight in the 4 days post infection compared to NK-cell sufficient mice (Fig. 2E, male, and Fig. 2F, female). While unvaccinated mice reached their humane endpoint of 20% weight loss by 6 days post infection, vaccinated animals lost only 5% of their body weight and recovered to pre-infection weights by day 8 (Fig. 2G). However, vaccinated and infected animals which lacked NK cells had prolonged weight loss which was more severe (10%) than in NK cell-intact vaccinated mice (5%) and recovered to baseline only by day 14 (Fig. 2G).

### Reduction of IFN-γ with influenza and NK cell depletion

Given that vaccinated NK cell-depleted animals lost more weight than vaccinated NK cell-intact animals despite a lower viral burden (Fig. 2), we next looked at inflammatory markers in the lung and plasma. In agreement with data from lung supernatants (Fig. 2D), viral burden determined from whole lung RNA preparations was significantly lower after NK cell depletion in unvaccinated animals (p=0.002), and not in vaccinated animals (p=0.08) (Fig. 3A). While *Il6* and *Ifnγ* expression were increased with influenza infection (Fig. 3B), regardless of vaccination status and viral burden (Fig. 3A), transcripts of *Ifnγ* were lower in NK cell depleted mice than in NK-intact mice (naïve p=0.004, vaccinated p=0.057).

**Figure 3.**
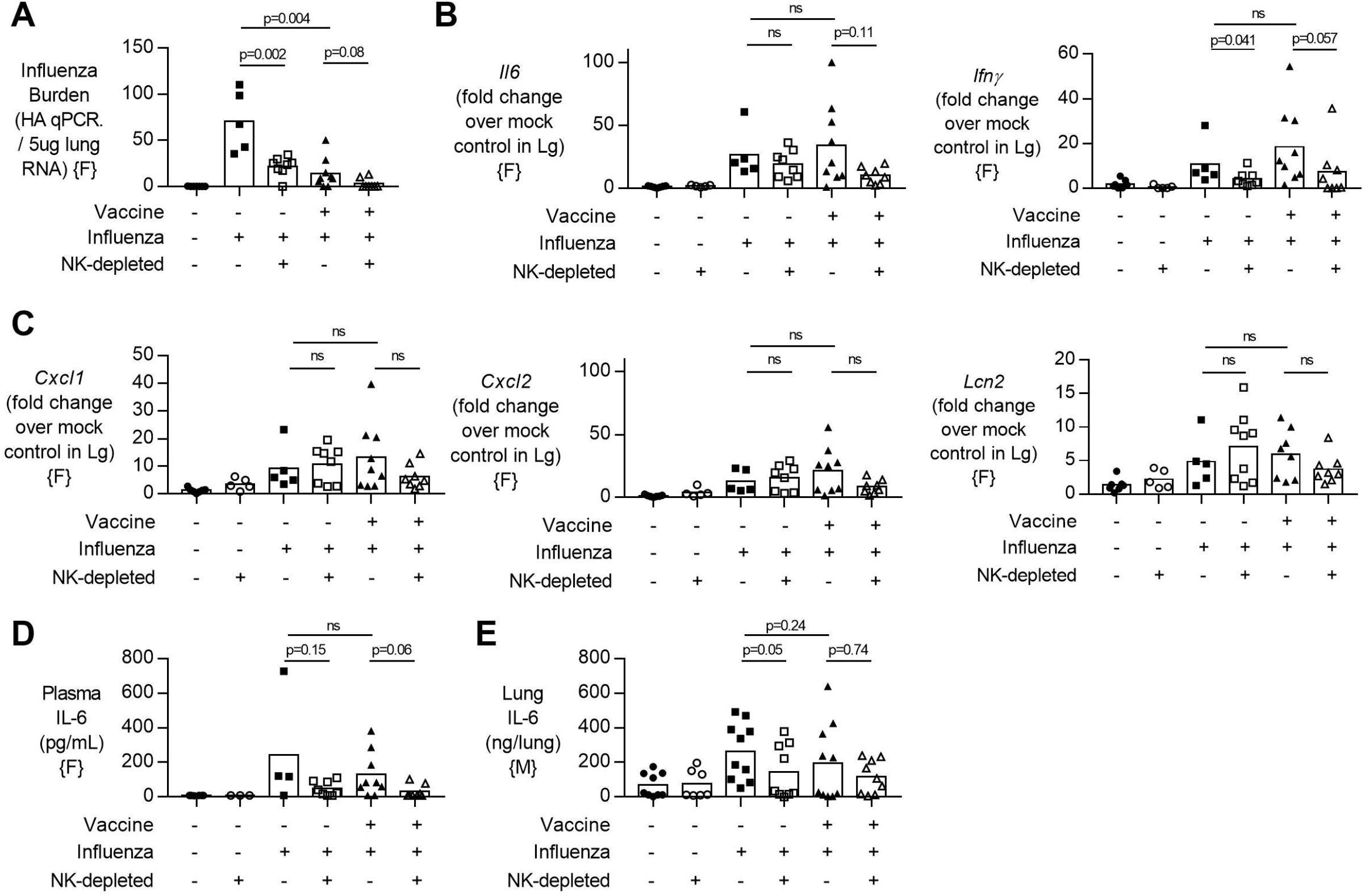
NK cell depletion reduces lung viral burden and lung IFN-γ in vaccinated, influenza-challenged mice. Four days post infection in the model described in Fig. 2A, **(A)** Lung RNA was analyzed by qPCR for influenza viral burden (plotted against a dose curve of IFA with known HAU, giving HAU equivalents per 5ug RNA tested). **(B-C)** Transcript levels of inflammatory cytokine genes **(B)** *Il6* and *Ifnγ* and **(C)** neutrophil-related chemokines *Cxcl1* and *Cxcl2*, along with neutrophil lipocalin protein *(Lcn2)* RNA induction normalized to housekeeping gene ß-actin and displayed as induction over mock-treated control mice. **(D)** Plasma levels of IL-6 (pg/mL). **(E)** IL-6 levels in lung supernatants after whole lung enzymatic digestion for single cell isolation and viral burden quantification. (A-E) A pool of two independent experiments (n=5-9/group; (A-D) were female {F} and (E) were male {E} mice used for single cell isolation). Dots represent individual mice with bars showing mean. Significance determined by Mann-Whitney U test, ns = not significant. Spearman correlation coefficient shown.

Given that neutrophil influx into an influenza-infected lung can enhance viral control (Dienz et al., 2012), but can also contribute to tissue damage through the release of extracellular traps (Narasaraju et al., 2011), we wondered whether increased neutrophil activity in NK cell-depleted mice might explain our findings. We therefore determined transcript levels of neutrophil chemokines (*Cxcl1*, *Cxcl2*) and neutrophil secreted protein *Lcn2* but found no significant differences in transcript concentrations between NK cell-deficient and NK cell-intact animals (Fig. 3C). Lastly, we assessed circulating levels of IL-6 (which have been implicated in neutrophil-mediated effects (Dienz et al., 2012)) and found a tendency towards lower plasma IL-6 concentrations in both vaccinated and unvaccinated NK-depleted mice compared to NK cell-intact mice (vaccinated p=0.15, unvaccinated p=0.06) (Fig. 3D), reflecting lower transcript levels in the lung (Fig. 3B). At the protein level in whole lungs after enzymatic digestion, IL-6 is lower in unvaccinated NK-depleted mice but not in vaccinated mice (vaccinated p=0.74, unvaccinated p=0.05) (Fig. 3E).

### Influenza-mediated lung pathology in vaccinated and NK cell-depleted mice

Given the decreased viral burden but increased weight loss upon influenza challenge in vaccinated NK cell-depleted mice compared to NK cell-sufficient mice (Fig. 2), we examined the lungs for histological evidence of pathology. H&E stained lung sections were assessed for pulmonary inflammation (vasculitis, bronchiolitis, and alveolitis), edema (perivascular, peribronchiolar, and alveolar), and infiltrating lymphocytes and neutrophils (perivascular, peribronchiolar, and alveolar) (Fig. 4; see Supplementary Table 1 for individual mouse data). Influenza infection alone induces severe pathology (Fig. 4). Reduced viral load after vaccination was accompanied by reduced pulmonary leukocyte influx (p=0.024), with a trend for reduced pulmonary inflammation (p=0.065) and edema (p=0.09). NK cell depletion *per se* (i.e. in the absence of infection) induced mild lung pathology with evidence of inflammation (p=0.008) and leukocyte infiltration (p=0.006). However, after influenza infection, pathology was significantly less severe in NK-depleted mice compared to NK cell-sufficient mice (inflammation p=0.04, leukocyte infiltration p=0.048). Interestingly, while weight loss was more severe in vaccinated and influenza challenged NK-depleted mice than in NK cell-sufficient mice, pathological changes in the lungs were not significantly different between the two groups of mice (Fig. 4).

**Figure 4.**
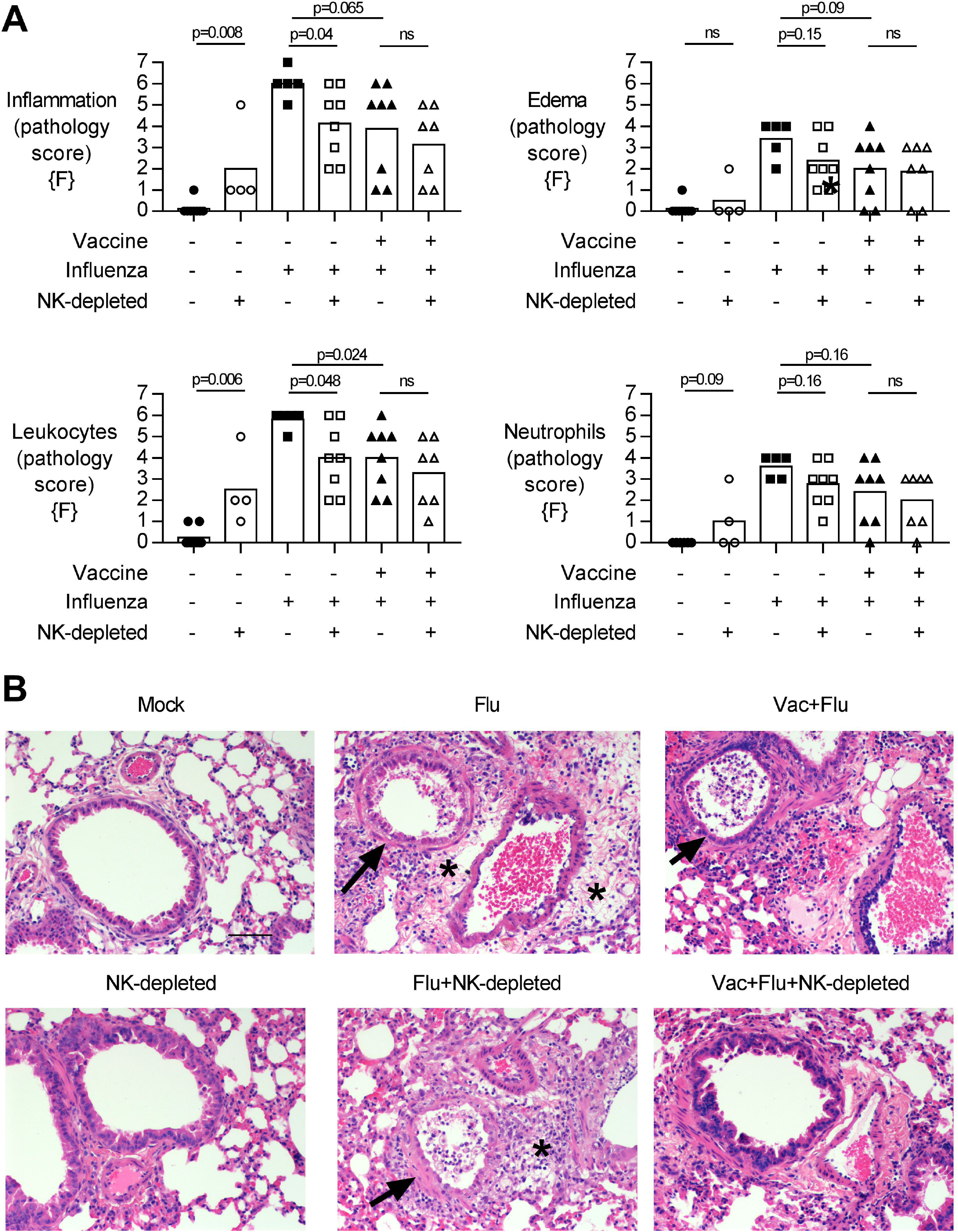
NK cell depletion in vaccinated, influenza-challenged mice does not alter pathology. Four days post infection in the model described in Fig. 2A, whole lungs were excised, stored in 10% formalin, and embedded on paraffin for hematoxylin and eosin staining. Pathology was scored for: **(A)** Inflammation (vasculitis, bronchiolitis, and alveolitis), Edema (perivascular, peribronchiolar, and alveolar), Leukocytes and Neutrophils (in perivascular space, peribronchiolar space, and alveolar wall). Full scoring details in supplementary files. Pathology scored from a pool of two independent experiments (n=5-9/group); all female {F} mice. Dots represent individual mice with bars showing mean. Significance determined by Mann-Whitney test, ns = not significant. **(B)** Representative photomicrographs of pathological changes observed. Arrows, bronchiolitis with exudates in bronchiolar lumina. * Perivascular and peribronchiolar edema. Scale bar in mock equals 50 μm.

Finally, we quantified infiltrating leukocytes and neutrophils in the lung by flow cytometry (Fig. 5). Influenza infection led to an infiltration of activated (CD69+) CD3+ T cells into the lung; this was not affected by NK-depletion but was attenuated in vaccinated mice (Fig. 5A). There were no obvious effects of infection or vaccination on CD19+ B cell populations in the lung; the modest apparent increase in B cell proportions in NK cell-depleted mice may simply reflect distortion of cell percentages by the absence of NK cells (Fig 5B). However, given that influenza infection increases the influx of CD11b+ cells into the lung (Fig. 1D), we further characterized influx of both inflammatory monocytes (%Ly6C-high) and neutrophils (%Ly6G+) in the lungs of NK-depleted mice. The modest reduction in leukocyte infiltration in the lung that was evident from histology (Fig. 4), was reflected in reduced infiltration of inflammatory monocytes into the lung in vaccinated NK cell-sufficient animals compared to unvaccinated animals (p=0.005) and in infected NK cell-depleted animals compared to NK cell intact animals (p=0.002) (Fig. 5C) but there was no additional effect of combining vaccination and NK cell depletion (p=0.99). Neutrophil infiltration into the lung was also reduced by vaccination in NK cell intact mice (p = 0.007) but in this case, the effect was reversed by NK cell depletion such that neutrophil infiltration was higher in vaccinated NK cell-depleted mice than in vaccinated NK cell intact mice, p=0.03 for proportion of neutrophils (Fig. 5D) and p=0.06 for absolute counts (data not shown), and correlated with weight loss (Fig. 5E). Taken together, these data suggest that the increased weight loss observed with influenza infection in NK cell-depleted vaccinated mice compared to NK cell intact vaccinated mice may be due, in part, to failure of NK cell-depleted mice to control neutrophil infiltration into the lungs, despite reduced viral burden.

**Figure 5.**
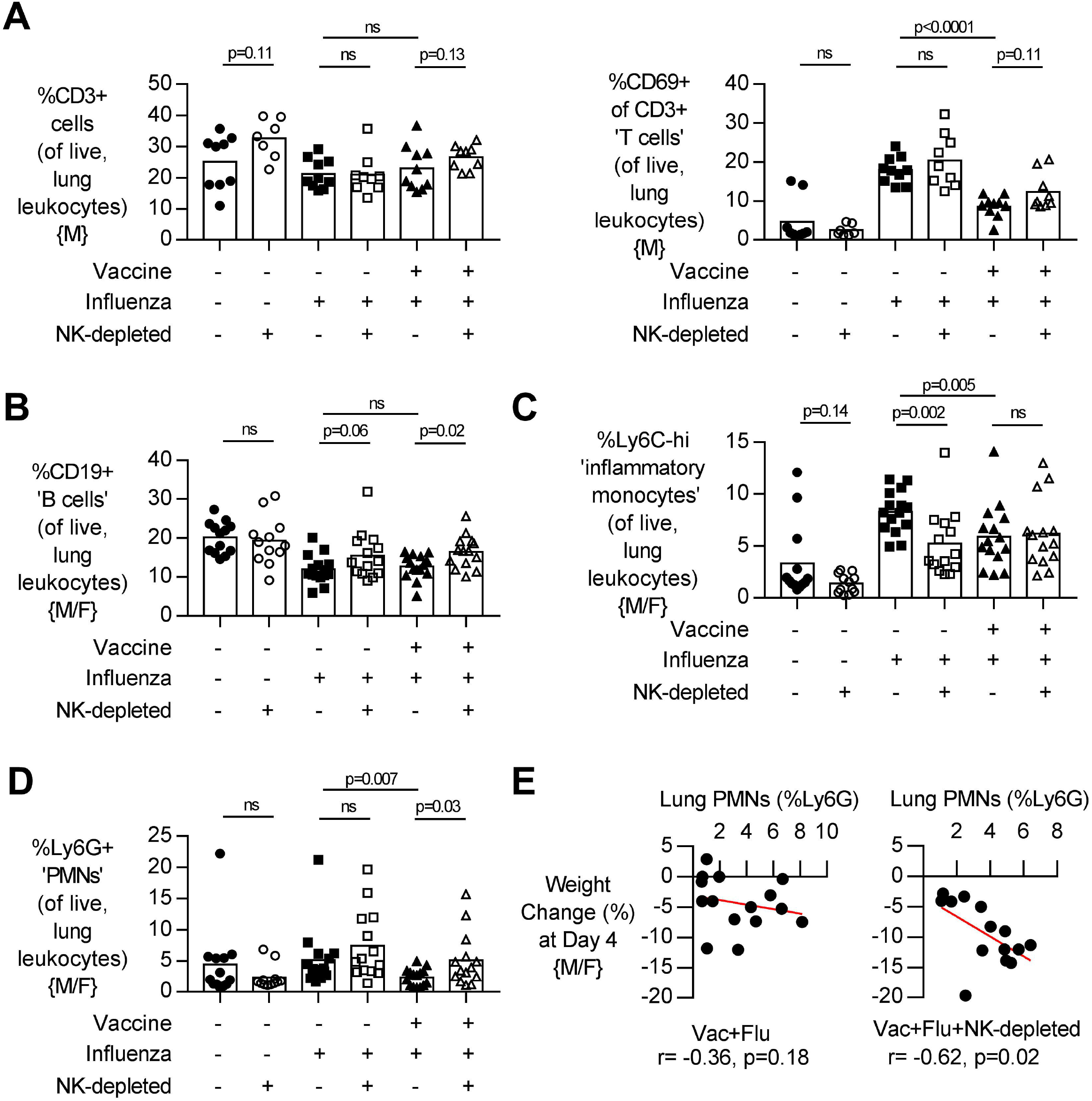
NK cell depletion in vaccinated, influenza-challenged mice increases lung neutrophil infiltration. Four days post infection in the model described in Fig. 2A, whole lungs were excised and single cells isolated for flow cytometry. **(A)** Proportion (%) of CD3+ T cells and active (CD69+) CD3+ T cells, as determined from singlet, live lung leukocytes. Proportion (%) of **(B)** CD19+ B cells, **(C)** Ly6C-high inflammatory monocytes, and **(D)** Ly6G+ neutrophils, with neutrophils plotted against weight loss **(E)**. (A) T cell data from a pool of two independent experiments (n=7-10/group); all mice were male {M}. (B-E) Data a pool of three independent experiments (n=7-10/group of male {M} mice) and (n=4-5/group of female {F} mice). Dots represent individual mice, with bar representing mean. Significance determined by Mann-Whitney U test, ns = not significant. Spearman correlation coefficient and p-value shown.

### Depletion of NK cells after vaccination, and subsequent repopulation, does not alter the response to influenza challenge in mice

To determine if the effects of NK cell depletion on post vaccination immunity to influenza were mediated by NK cells present at (and potentially affected by) vaccination (Goodier et al., 2016), we depleted NK cells (by a single treatment with 2.5 μg of DT) 3 weeks after influenza vaccination (as previously) but then waited another 3 weeks before challenging the mice (to allow repopulation) (Fig. 6A). NK cells were initially (after 3 days) very effectively depleted by the DT treatment and, as predicted, the NK cell compartment did partially recover by the time of influenza challenge (Fig. 6B). In this case, removal of NK cells present at vaccination and subsequent NK cell repopulation resulted in influenza infections that were not significantly different from those in intact mice, with no significant differences in disease severity (weight loss; Fig. 6C), lung viral load (Fig. 6D) or circulating IL-6 concentrations (Fig. 6E). This experiment suggests that the effects of NK cell depletion on post vaccination immunity are due to the lack of all NK cells, rather than a lack of NK cell populations that were primed or activated by vaccination.

**Figure 6.**
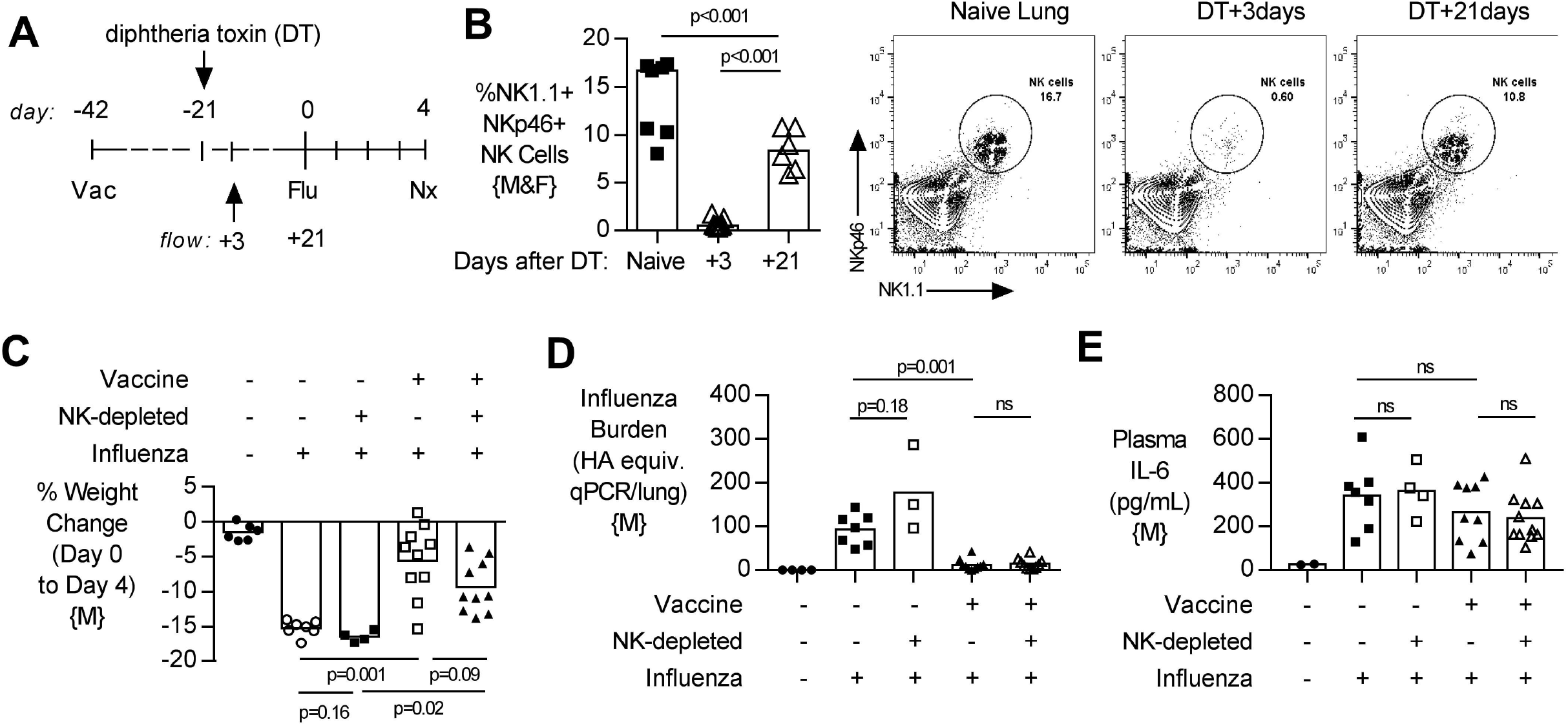
Depletion of NK cells after vaccination, and subsequent repopulation, does not alter lung viral burden, disease severity or systemic inflammation after challenge. **(A)** Transgenic C57BL/6 mice with NKp46 driven expression of diphtheria toxin (DT) receptor were vaccinated 42 days (d) prior to intranasal influenza (Flu) challenge and treated with DT (NK-depleted) 21 days prior to challenge with necropsy (nx) at 4 days post influenza challenge. **(B)** At 3 and 21 days post DT treatment, lungs were excised and single cells isolated for flow cytometry for the proportion (%) of NK1.1+, NKp46+ NK cells. **(C)** Weight loss at 4 days post influenza challenge. **(D)** Lung cell-free supernatants were analyzed by qPCR for influenza viral burden (plotted against a dose curve of Flu with known HAU, giving HAU equivalents). **(E)** Plasma levels of IL-6 (pg/mL). (B) NK cell depletion data a pool of two independent experiments, (n=2/group of male {M} mice) and (n=5-6/group of female {F} mice). (C-D) Data a pool of two independent experiment (n=4-10/group); all male {M} mice. Dots represent individual mice with bars showing mean. Significance determined by Mann-Whitney test, ns = not significant.

## Discussion

The key findings of this study, summarized in Fig. 7, are that: (1) influenza vaccination is effective in reducing viral burden and weight loss in mice; (2) in both vaccinated and unvaccinated mice, NK cells ameliorate disease (weight loss) at the expense of delaying viral clearance; (3) that the magnitude of the effect on disease is greater in vaccinated than unvaccinated mice (2.5-fold greater weight loss vs 1.11-fold, respectively), and (4) the depletion of any ‘memory’ NK cells that might have been induced by vaccination did not alter the response to influenza virus in vaccinated mice. These data suggest that NK cells play an important homeostatic role, allowing influenza virus to be controlled without causing severe disease, and that this effect is enhanced by vaccination – indicating a role for NK cells in regulating the adaptive immune response. Our study therefore supports previous data suggesting an important role for NK cells in moderating adaptive immune effector mechanisms (Pallmer and Oxenius, 2016).

**Figure 7.**
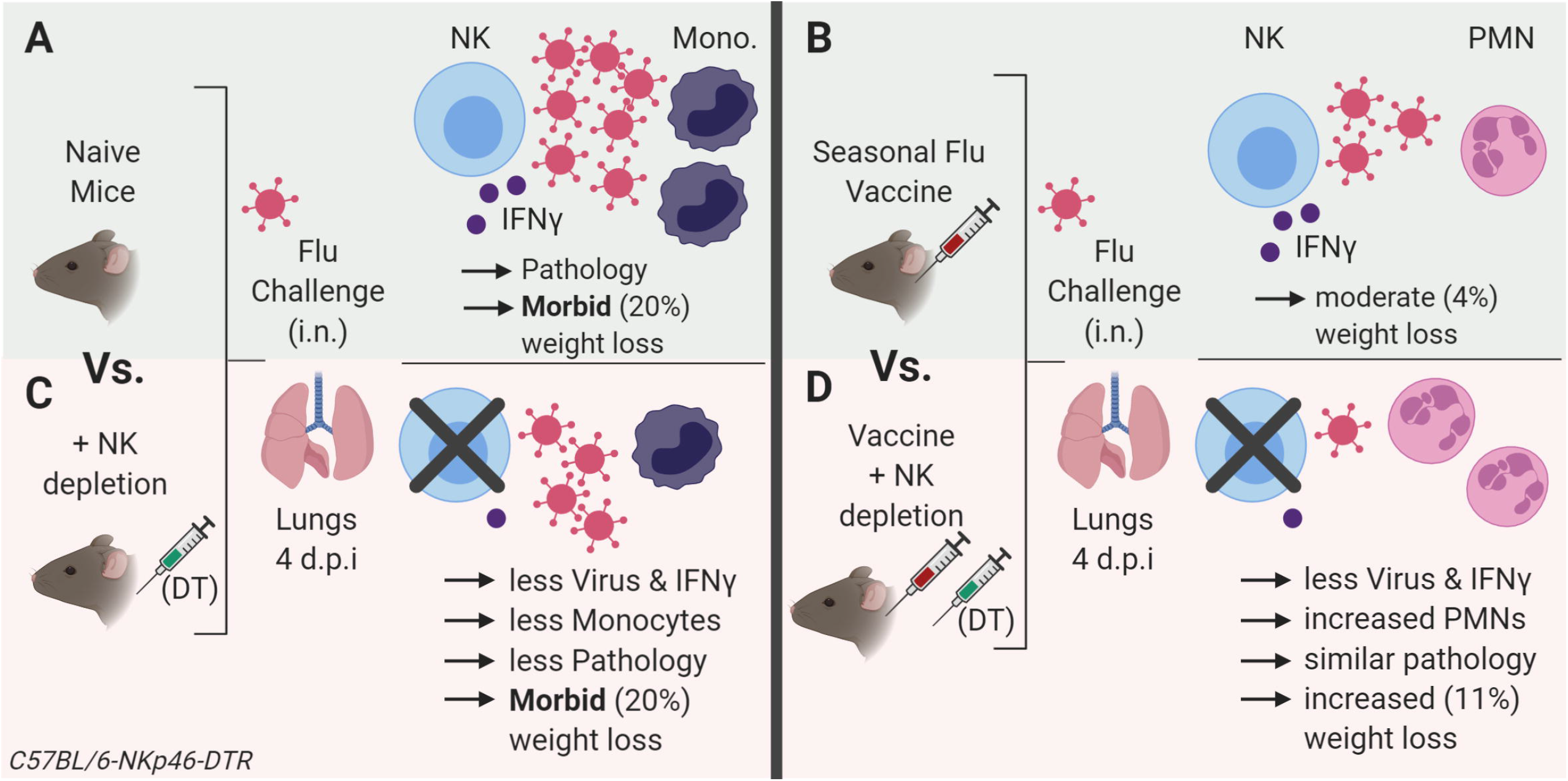
Summarized model of changes seen with influenza vaccination and challenge with NK cell depletion. **(A)** Influenza virus infection in naïve mice results in robust lung pathology, characterized by interferon (IFN)-γ mRNA and inflammatory monocytes (%Ly6C-high) influx with rapid weight loss at 4 days post infection. **(B)** Vaccination with the human Sanofi trivalent influenza vaccine reduces lung viral burden, pathology and monocytes with minor weight loss (4%). **(C)** In unvaccinated, naïve animals which were infected with influenza, NK cell depletion reduced viral burden, IFN-γ mRNA, pathology, and monocyte influx. However, these mice still progressed to 20% weight loss by day 4 post infection. **(D)** In vaccinated mice which were lacking NK cells, influenza virus burden was reduced even further and robust IFN-γ from vaccination now lower, yet total pathology score was similar to NK intact mice – however with PMNs now increased in frequency in the lung by flow cytometry. Despite a reduced viral burden, these animals showed sustained weight loss at 11%. Model created with BioRender.

Previous work has suggested that the role of NK cells during influenza infection is dependent on the infecting dose of the virus, with NK cells being protective during low dose infection (0.5 HAU) and pathogenic during high dose infection (5 HAU) (Zhou et al., 2013). Although we used the A/California/4/2009 strain of influenza rather than PR8 strain used by Zhou *et al*, we used an infecting dose that is similar to the low dose used by Zhou et al. (0.5 HAU) and saw a similar trend. Interestingly, an opposite effect was seen during murine cytomegalovirus (MCMV) infection where NK cells prevented pathology at high viral doses but enhanced disease at low doses (Waggoner et al., 2011). In our study, NK cell-depleted unvaccinated male mice lost significantly more weight than intact mice after infection (Fig. 2E); however this effect was not seen in female mice (Fig. 2F) suggesting that NK cells moderate disease severity differently in male and female mice despite the two sexes having similar viral loads. Regardless, in NK-dell depleted and vaccinated mice we saw significant weight loss in both sexes (Fig. 2E and 2F), suggesting that sex differences are ablated by vaccination.

The mechanisms by which NK cells moderate adaptive immune responses are not fully understood. Our observation that NK cell depletion tended to increase neutrophil influx into the lungs of influenza infected mice, and significantly increased neutrophil accumulation in lungs of vaccinated mice, suggests that NK cells may serve to limit neutrophil migration to sites of infection. This would be in line with studies showing that neutrophils are important in controlling influenza virus infection (Tate et al., 2008;Tate et al., 2009;Tate et al., 2011) but can contribute to severe pulmonary pathology (Tumpey et al., 2005;Perrone et al., 2008;Feng et al., 2015).

Another possibility is that NK cells regulate the accumulation of adaptive, cytotoxic (CD8+) effector T cells at the site of infection, thereby reducing tissue damage but slowing viral clearance. In RSV infection, for example, CD8+ cells are directly associated with weight loss and depleting them reduces disease (Tregoning et al., 2008). NK cells have been shown to eliminate activated CD4+ and CD8+ T cells in different model systems (Waggoner et al., 2011;Lang et al., 2012;Pallmer and Oxenius, 2016), mediated by either perforin or TNF-related apoptosis-inducing ligand (TRAIL) (Peppa et al., 2013;Schuster et al., 2014). However, thus far, in influenza vaccination models CD8+ T cells appear to contribute to both reduced viral load and reduced disease severity (Lambert et al., 2016). Unfortunately, we did not determine CD8 T cell responses in this current study and therefore further work is needed to determine whether the lack of NK cells removes the brake on adaptive T cell responses during influenza infection.

In a recent study (Zamora et al., 2017), murine NK cells licensed on self MHC were shown to localize to infected lung tissue and produce IFN-γ after influenza A (strain PR8) infection. Unlicensed NK cells, in contrast, were enriched in draining (mediastinal) lymph nodes, produced GM-CSF and promoted dendritic cell infiltration and CD8+ T cell responses. It is therefore likely that distinct subsets of NK cells may selectively promote inflammation or antiviral immunity. Furthermore, this may differ depending on levels of adaptive immunity.

In humans, memory-like NK cells can be generated by cytokine or influenza virus pre-activation; these cells show enhanced responses to cytokines or influenza virus upon re-stimulation (O’Sullivan et al., 2015;Goodier et al., 2016;Wagstaffe et al., 2018). It is conceivable that prior activation of NK cells, for example by vaccine-induced inflammatory cytokines, could influence the migration and function of these cells upon viral challenge. Certainly, human nasal challenge with Fluenz induces a local (nasal mucosal) innate cytokine signature with potential NK cell activating capacity (Jochems et al., 2018). Nevertheless, we wondered whether a similar phenomenon might also be apparent in this mouse model (where more precise dissection of the underlying local mechanism of NK cell enhancement might be possible). Therefore, we depleted NK cells immediately after vaccination (thus removing any putative ‘memory’ NK cells, defined as cells with altered function after prior exposure), and rested the mice for three weeks to allow repopulation before challenging with influenza virus. We observed no differences in the outcome of influenza infection between these NK cell-depleted/repopulated vaccinated mice and intact, vaccinated mice. These data suggest that “memory” NK cells are not induced by influenza vaccination in mice or that any such cells are no more effective at moderating adaptive responses to influenza than are “naïve” NK cells. In mice, cytokine-induced memory-like NK cells can be adoptively transferred and maintained by homeostatic proliferation (Keppel et al., 2013), but have to date not been demonstrated after influenza vaccination. Human studies have been limited to peripheral blood and further investigations are needed to test how NK cells influence inflammatory cellular infiltrates at the site of virus infection. Vaccination and challenge studies (with local mucosal sampling for virus, cellular infiltrates and cytokine production) are now needed to reveal the impact of vaccine-induced NK cell priming and “memory” generation on virus replication and pathology.

There are some important caveats to keep in mind, however. Firstly, in contrast to previous studies using mice lacking NK cells at the time of infection or vaccination, or depleted of NK cells with antibodies on animals of different sexes (Table 1), we used an inducible, endogenous method of NK cell depletion which may have fewer limitations (Walzer et al., 2007). Nevertheless, DT treatment alone lead to some limited weight loss and pulmonary pathology, likely due to death of approx. 15% of lung leukocytes, although there was no induction of inflammatory cytokines nor any influx of immune cells into the lung. It remains a possibility, therefore, that the mild inflammation induced by NK cell depletion may have affected the outcome of the experiments. Further, sex differences in influenza-mediated lung pathology have been noted (Klein et al., 2012). Secondly, the ethically-approved humane end point for our study was deemed to be 20% weight loss (after which animals were euthanized), essentially precluding determination of long-term survival after NK cell depletion but some studies have shown only limited correlation between weight loss and long term survival (Abdul-Careem et al., 2012), with viral challenge dose a greater predictor of survival despite similar weight loss kinetics (Zhou et al., 2013). A further caveat of the Rosa-DT conditional ‘NK cell’ depletion system used here is a potential parallel depletion of mature NKp46+ ILC1 and ILC3 cells (Cortez et al., 2015). Studies in deficient *ncr1*^gfp^ mice, which are susceptible to lethal influenza A infection, demonstrate systemic loss of ILC1, whilst maintaining NKp46 negative NK cell subsets (Gazit et al., 2006;Wang et al., 2018). Although rare in the non-pathologic lung, our studies do therefore not exclude a potential contribution of ILC-1s to inflammatory processes after influenza vaccination or challenge infection. Lastly, characterization of cytokine protein levels in the bronchoalveolar lavage fluid would be beneficial in understanding transcriptional changes observed in this study.

To summarize, we have demonstrated that NK cells play a homeostatic role in adaptive immunity to influenza infection in mice. While the precise mechanism by which NK cells modulate adaptive immunity remains unclear, their presence is crucial for resolving infection with minimal immune pathology. Further studies to determine the mechanism(s) at play may inform the design of safer and more effective influenza vaccines.

## Supporting information

Supplemental Flow Cytometry Gates

Supplemental Pathology Scores

## Acknowledgements

We thank Professor Eric Vivier (Aix-Marseille, France) for generously supplying breeding pairs of C57BL/6J-NKp46:iCre^+/+^ mice, the LSHTM animal care staff for their technical assistance, John McCauley and Andreas Wack (Francis Crick Institute) for advice on the mouse influenza model and Christian Bottomley (LSHTM) for statistical advice.

## Author Contributions Statement

Study concept and design: JM, MG, and ER; data generation and analysis: JM, TQ, MK, AP, HG; drafting and revision of manuscript: JM, TQ, MK, AP, JT, MG, and ER; critical appraisal and approval for submission: all authors.

## Data Availability Statement

The raw data supporting the conclusions of this manuscript will be made available by the authors, without undue reservation, to any qualified researcher.

