## Supplemental Flow Cytometry Gates for "Natural Killer cells dampen the pathogenic features of recall responses to influenza infection"

**Supplementary Digital Content:**

Supplementary Figure Legends:

**Figure S1, related to Figure 1D.** Representative flow cytometry gating strategy for CD19+ B cells and CD3+ T cells. C57BL/6J lung cells were isolated as described in methods. Gating was performed by singlet (FSC-A v FSC-H), leukocytes (removing high SSC-A and low SSC-A/FSC-A), live cells. Values shown as frequency (%) of parent population.

**Figure S2, related to Figure 1D-E.** Representative flow cytometry gating strategy for NKp46+ NK1.1+ Natural Killer (NK) cells. C57BL/6J lung cells were isolated as described in methods. Gating was performed by singlet (FSC-A v FSC-H), leukocytes (removing high SSC-A and low SSC-A/FSC-A), live cells. Values shown as frequency (%) of parent population.

**Figure S3, related to Figure 2B.** Representative flow cytometry gating strategy for depletion of NKp46+ NK1.1+ Natural Killer (NK) cells with diphtheria toxin (DT). NKp46-DTR lung cells were isolated as described in methods. Gating was performed by singlet (FSC-A v FSC-H), leukocytes (removing high SSC-A and low SSC-A/FSC-A), live cells. Values shown as frequency (%) of parent population

**Figure S4, related to Figure 5A.** Representative flow cytometry gating strategy for CD3+ T cells and their activation via CD69. NKp46-DTR lung cells were isolated as described in methods. Gating was performed by singlet (FSC-A v FSC-H), live cells, and leukocytes (removing high SSC-A and low SSC-A/FSC-A). Values shown as frequency (%) of parent population

**Figure S5, related to Figure 5B-D.** Representative flow cytometry gating strategy for CD19+ B cells, Ly6C-high (hi) 'inflammatory' monocytes, and Ly6G+ Neutrophils. NKp46-DTR lung cells were isolated as described in methods. Gating was performed by singlet (FSC-A v FSC-H), live cells, and leukocytes (removing high SSC-A and low SSC-A/FSC-A). Values shown as frequency (%) of parent population.

### Supplementary Figures:

Figure S1:

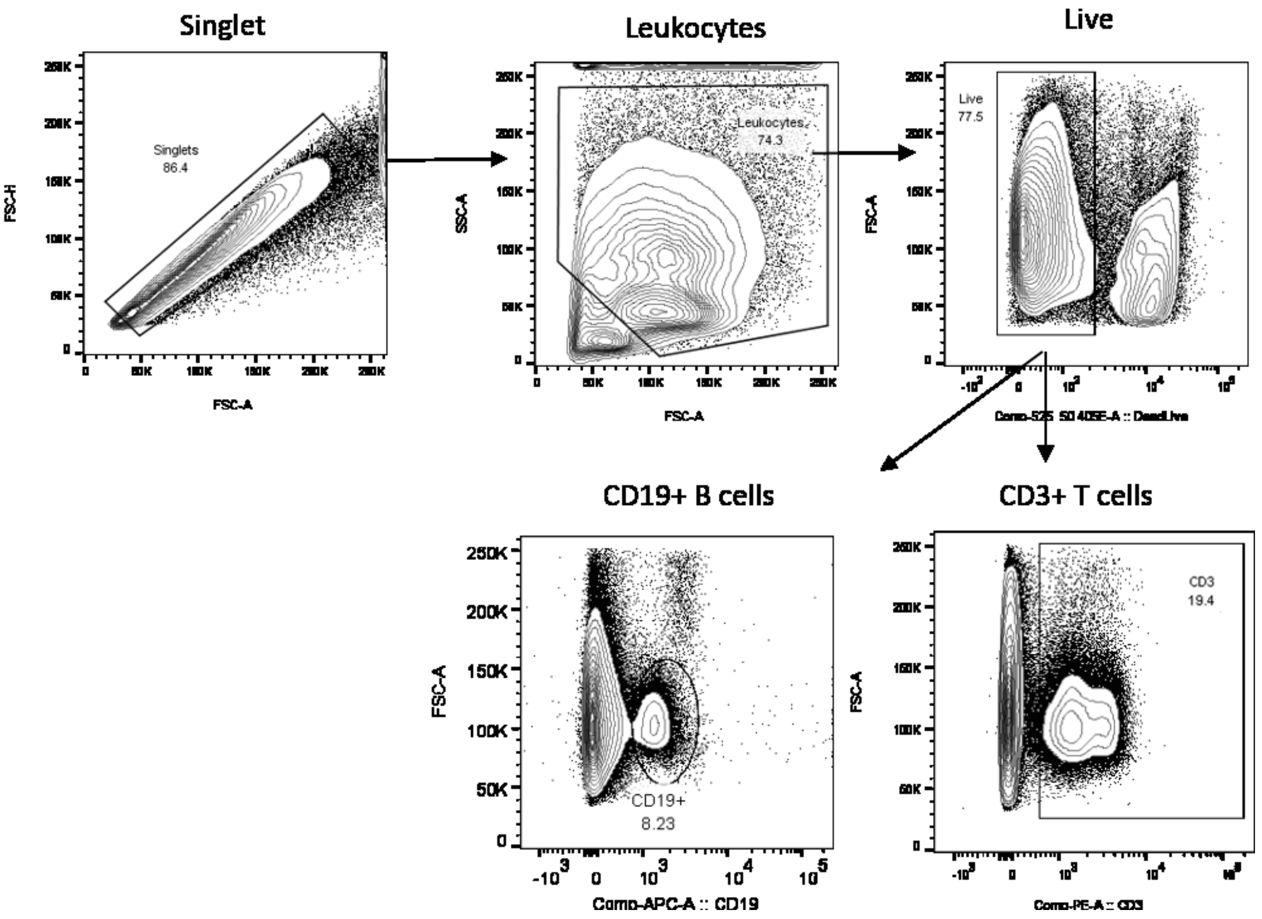

Figure S2:

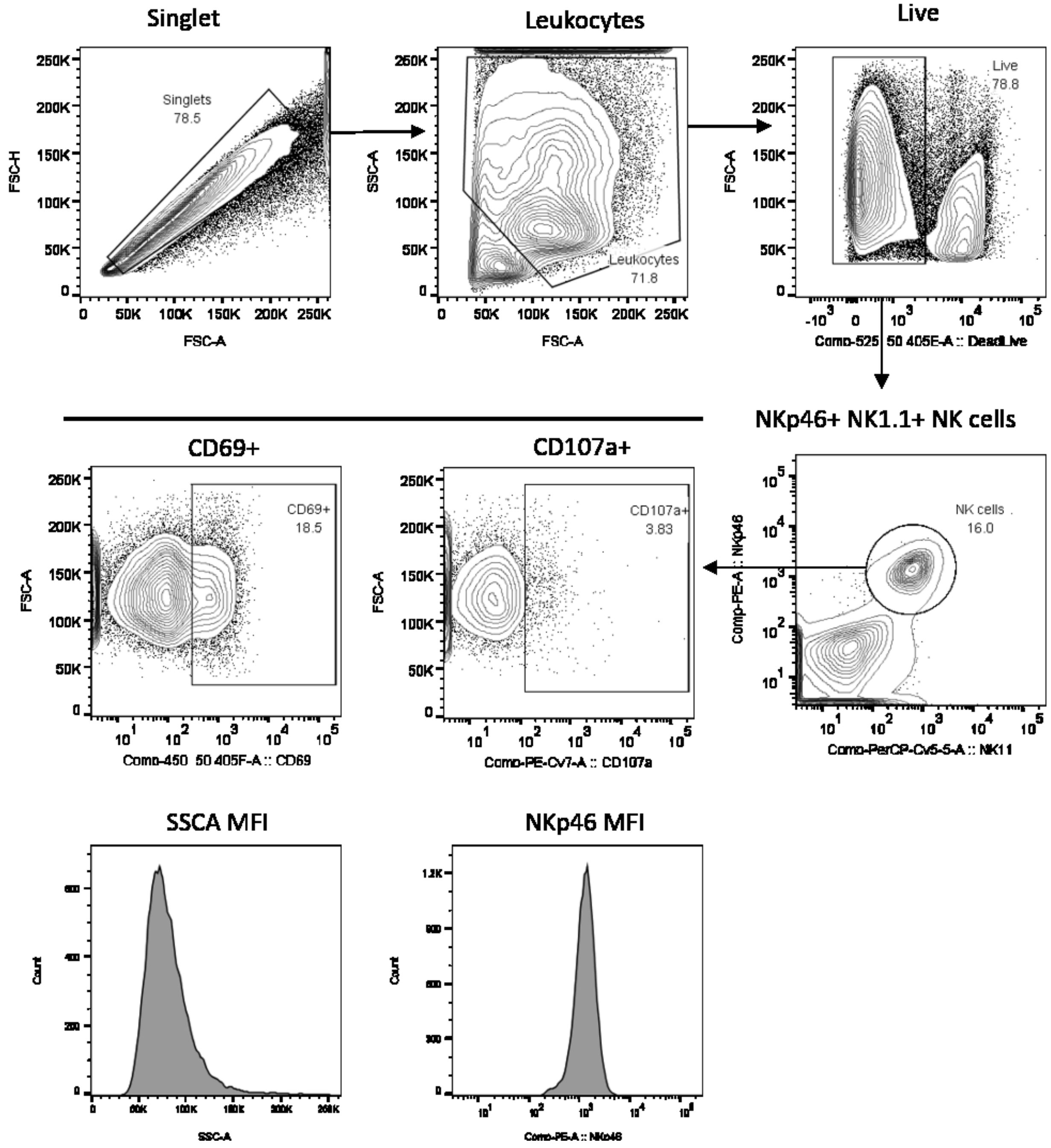
Singlet
Singlets
78.5
FSC-H
FSC-A
Leukocytes
Leukocytes
71.8
SSC-A
FSC-A
Live
Live
78.8
FSC-A
Comp-525_50 405E-A :: DeadLive
NKp46+ NK1.1+ NK cells
NK cells
16.0
Comp-PE-A :: NKp46
Comp-PerCP-Cy5-5-A :: NK11
CD69+
CD69+
18.5
FSC-A
Comp-450_50 405F-A :: CD69
CD107a+
CD107a+
3.83
FSC-A
Comp-PE-Cy7-A :: CD107a
SSCA MFI
Count
SSC-A
NKp46 MFI
Count
Comp-PE-A :: NKp46

Figure S3:

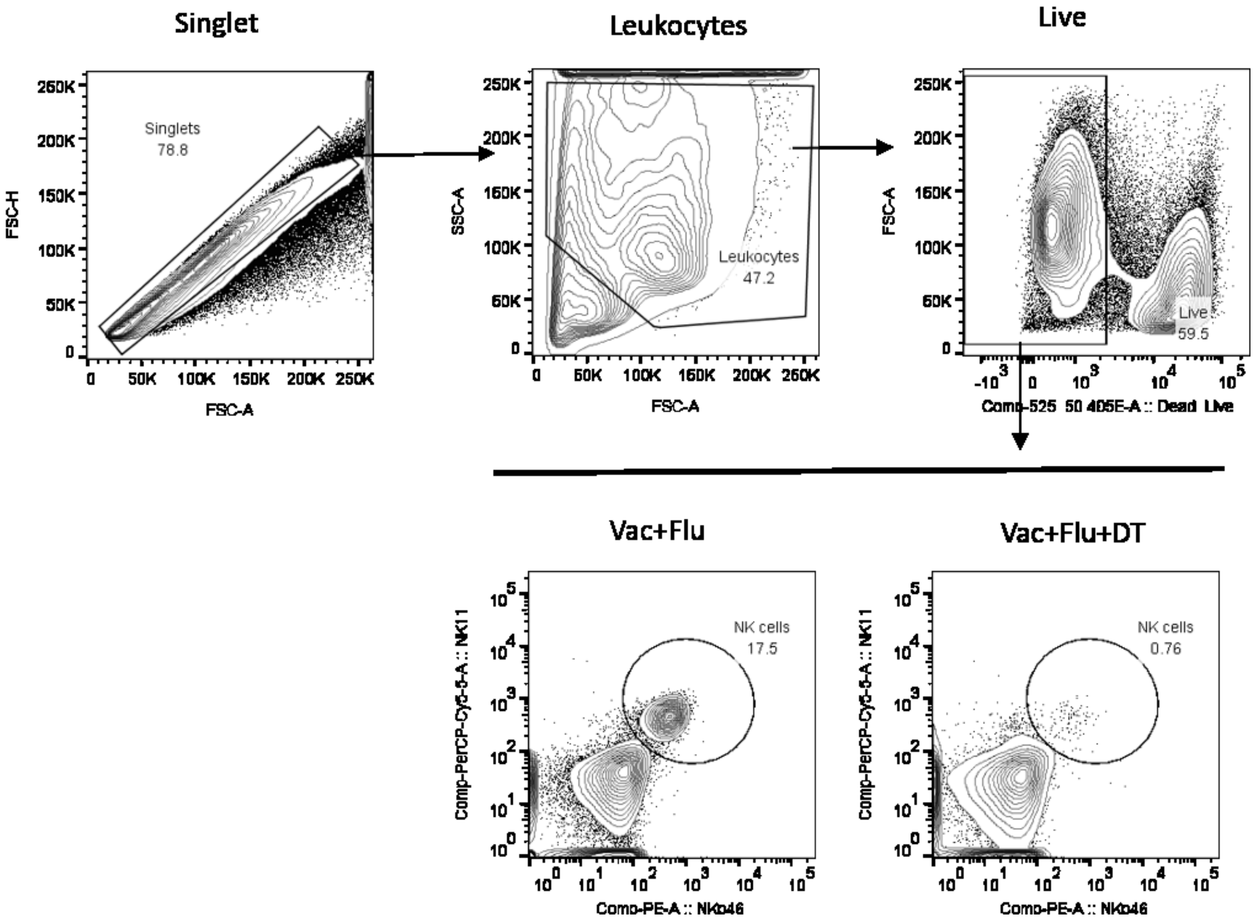
Singlet
Singlets
78.8
FSC-H
FSC-A
Leukocytes
Leukocytes
47.2
SSC-A
FSC-A
Live
Live
59.5
FSC-A
Comp-525 50 405E-A :: Dead Live
Vac+Flu
NK cells
17.5
Comp-PerCP-Cy5-5-A :: NK11
Comp-PE-A :: NKp46
Vac+Flu+DT
NK cells
0.76
Comp-PerCP-Cy5-5-A :: NK11
Comp-PE-A :: NKp46

Figure S4:

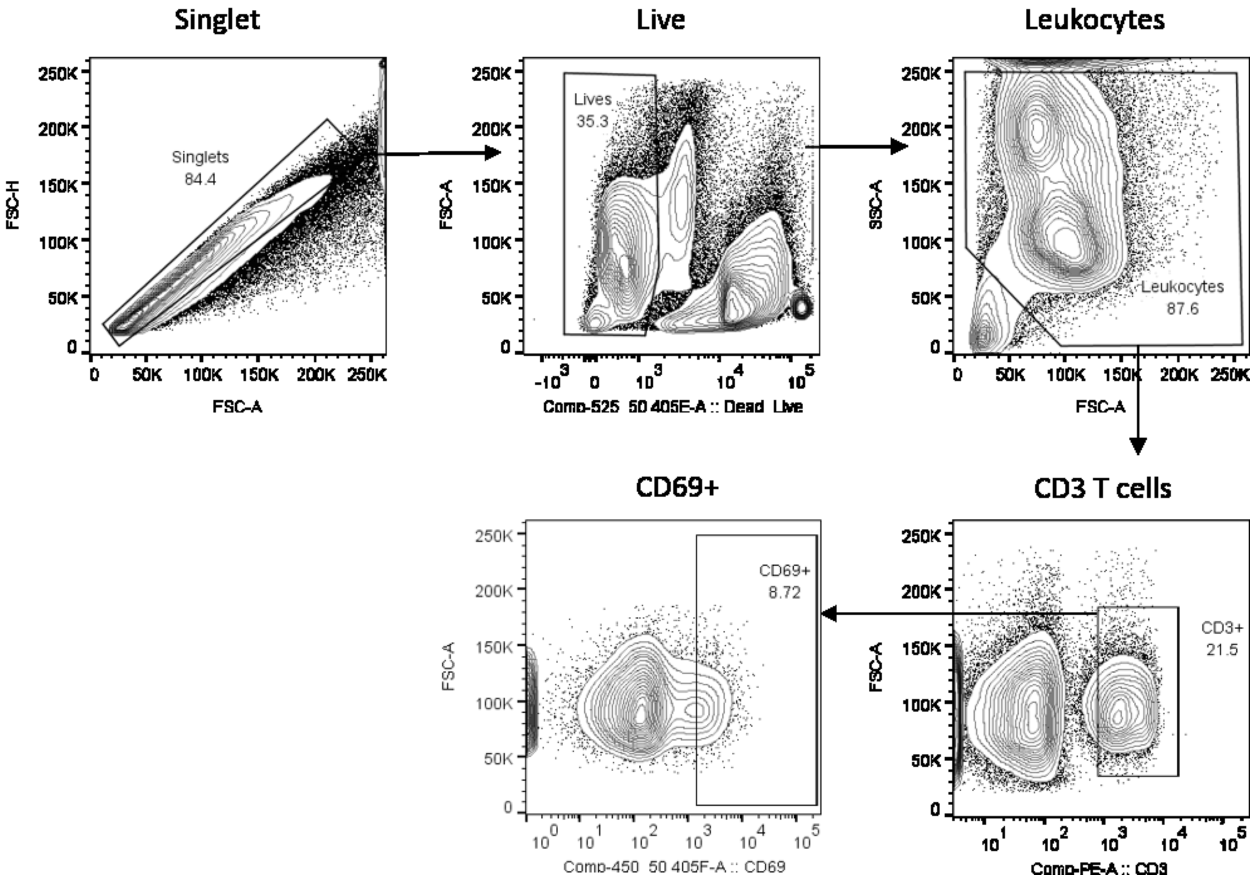

Singlet
Singlets
84.4
FSC-H
FSC-A
Live
Lives
35.3
FSC-A
Comp-525_50 405E-A :: Dead_Live
Leukocytes
Leukocytes
87.6
SSC-A
FSC-A
CD69+
CD69+
8.72
FSC-A
Comp-450_50 405F-A :: CD69
CD3 T cells
CD3+
21.5
FSC-A
Comp-PE-A :: CD3

Figure S5:

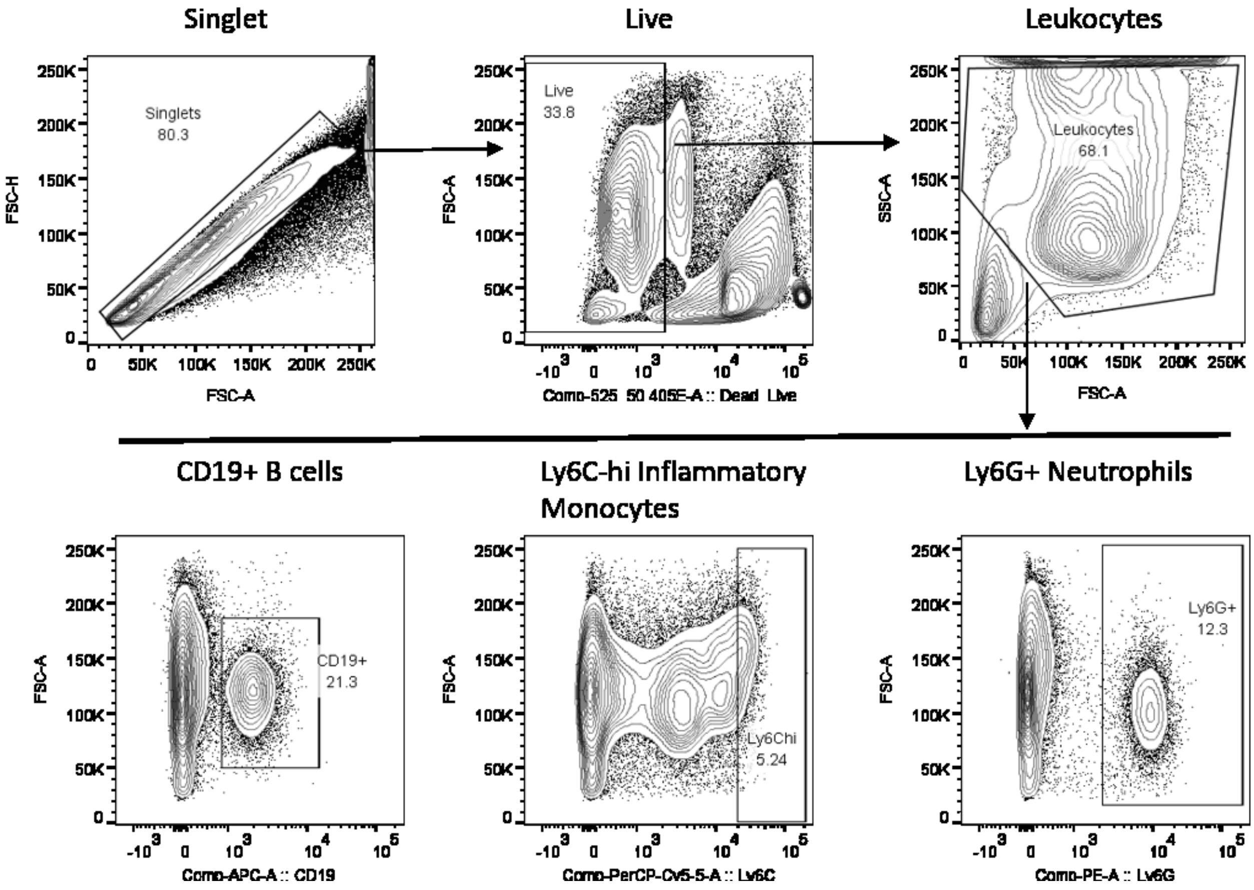
Singlet
Live
Leukocytes
Singlets
80.3
Live
33.8
Leukocytes
68.1
FSC-H
FSC-A
SSC-A
Comp-525_50 405E-A :: Dead_Live
CD19+ B cells
Ly6C-hi Inflammatory Monocytes
Ly6G+ Neutrophils
CD19+
21.3
Ly6Chi
5.24
Ly6G+
12.3
Comp-APC-A :: CD19
Comp-PerCP-Cy5-5-A :: Ly6C
Comp-PE-A :: Ly6G
